# The Liquid Structure of Elastin

**DOI:** 10.1101/095927

**Authors:** Sarah Rauscher, Régis Pomès

**Affiliations:** Molecular Structure and Function, The Hospital for Sick Children, Toronto, Ontario, Canada, M5G 0A4; Department of Biochemistry, University of Toronto, Toronto, Ontario, Canada, M5S 1A8

**Keywords:** elastin, molecular dynamics, intrinsically disordered proteins, protein aggregation, self-assembled elastomer, protein liquid-liquid phase separation, polymer melt

## Abstract

The protein elastin imparts extensibility, elastic recoil, and resilience to tissues including arterial walls, skin, lung alveoli, and the uterus. Elastin and elastin-like peptides are intrinsically disordered hydrophobic proteins that undergo liquid-liquid phase separation upon self-assembly. Despite extensive study for over eighty years, the structure of elastin remains controversial. We use molecular dynamics simulations on a massive scale to elucidate the structural ensemble of aggregated elastin-like peptides. Consistent with the entropic nature of elastic recoil, the aggregated state is stabilized both by the hydrophobic effect and by conformational entropy. The polypeptide backbone forms transient, sparse hydrogen-bonded turns and remains significantly hydrated even as self-assembly triples the extent of nonpolar side-chain contacts. The assembly approaches a maximally-disordered, melt-like state, which may be called the liquid state of proteins. These findings resolve long-standing controversies regarding elastin structure and function and afford insight of broad relevance to the phase separation of disordered proteins.

## INTRODUCTION

The elasticity of skin, lungs, major arteries, and other vertebrate tissues is imparted by elastin, a modular structural protein composed of alternating hydrophobic and cross-linking domains.^1^ While both types of domains contribute to the proper supramolecular assembly of the polymeric network, the cross-linking domains are understood to bestow cohesiveness and durability, whereas the hydrophobic domains are thought to confer the propensities for self-assembly and elastic recoil.^2^ Solutions of elastin and elastin-like peptides self-aggregate upon increasing temperature; this process, known as coacervation, is thought to be primarily due to self-association of hydrophobic domains. ^1^ Their capacity for temperature-controlled self-assembly make elastin-like peptides well-suited for biomaterials applications^3^ and drug delivery.^4^

Despite the biological importance of elastin and eighty years of study using a myriad of biophysical techniques,^5^ neither the molecular basis of self-assembly nor the structure of the self-assembled state are known. Numerous structural models of elastin have been proposed, which span a range from highly-ordered^6^ to maximally-disordered^7,8^ and emphasize either the hydrophobic effect^6,9—11^ or conformational entropy^7,8,12^ as the dominant contribution to the elastic recoil force.

High conformational entropy is the key feature of the earliest model proposed for elastin’s structure: the random network model.^7,8^ Based on thermoelasticity measurements indicating that elastin’s recoil force is almost entirely entropic, Hoeve and Flory postulated that elastin’s structure is an isotropic, rubber-like polymer network consisting of cross-linked, random chains.^7,8^ However, thermoelasticity measurements on elastin samples were carried out using alcohol diluents,^7,8,13,14^ and therefore rely on the assumption that alcohols do not significantly perturb the structure of elastin. The validity of this assumption has been questioned^9,15–17^ because the sequence composition of elastin is unusually enriched in hydrophobic residues, which likely interact with alcohols. Other types of mechanical studies have been carried out in a wide variety of solvents,^9,18^ including stress-strain measurements on elastin samples before and after glucose treatment.^19^ Whether or not the idealized random network model applies to the functional state of elastin is not known and remains controversial.

In contrast to the random network model, specific secondary structure preferences and significant burial of non-polar groups are common features of several other structural models of elastin.^6,9–11,16^ Elastin’s highly hydrophobic sequence led several groups to propose that the hydrophobic effect, rather than conformational entropy, is the major driving force of elastic recoil.^9–11,16^ Several models were proposed in which non-polar side chains are arranged to exclude water molecules; the most ordered of these models is the β-spiral, which consists of repeated β-turns.^6^ While the β-spiral model has been shown to be unstable,^11^ spectroscopic data are consistent with the presence ofβ-turns.^1,20^

To go beyond these largely qualitative and seemingly contradictory models, highresolution structural information is required. The conformational heterogeneity and self-association of elastin have impeded crystallographic and spectroscopic investigations and present a significant sampling challenge to molecular simulations.^21,22^ Accordingly, most previous computational studies of elastin-like peptides have been limited to molecular dynamics simulations of peptide monomers, starting with simulations of ~100 ps in vacuo^23–25^ and moving on to simulations of ~10-100 ns in explicit water.^11,19,22,26–31^ Although two of these studies examined peptide dimers^31^ or tetramers,^28^ to our knowledge simulations of larger aggregates of elastin-like (or, indeed, other intrinsically-disordered) peptides have never been reported. We have shown that attaining statistically-converged sampling of disordered elastin-like peptides necessitates simulation times in the microsecond time-range.^22,32^ To address these challenges, we performed large scale, all-atom molecular dynamics simulations of a monomer and an aggregate of 27 elastin-like peptides with a combined sampling time exceeding 200 *μ*s. As a result, we present the first atomistic description of the conformational ensemble of an elastin-like peptide successively in solution and in aggregated form.

## RESULTS AND DISCUSSION

### Peptide chain dimensions before and after self-assembly

We first compare the ensembles of the elastin-like peptide in solution (single chain, SC) and as an aggregate (multi-chain, MC) with respect to chain dimensions (Fig. 1). Both in solution and in the aggregate, the peptide chains sample heterogeneous, disordered structural ensembles, without a unique, preferred conformation. The ensembles differ significantly with respect to chain dimensions, with aggregated chains being much more expanded on average than the single chain in solution. Not only is the average radius of gyration, R_g_, higher for aggregated chains, the variance of R_g_ is also larger, indicative of a more heterogeneous underlying conformational ensemble. To quantify the difference in conformational entropy of the peptide chains in solution and in the aggregate, we compared the ensembles in terms of their RgD entropy, a metric recently introduced by Burger *et al.* to quantify the entropy of IDP ensembles.^33^ The RgD entropy of aggregated chains (2.254) is 25% higher than that of the solution ensemble (1.808) (Fig. S8), indicating a significant increase of conformational entropy upon aggregation.

**Figure 1.**
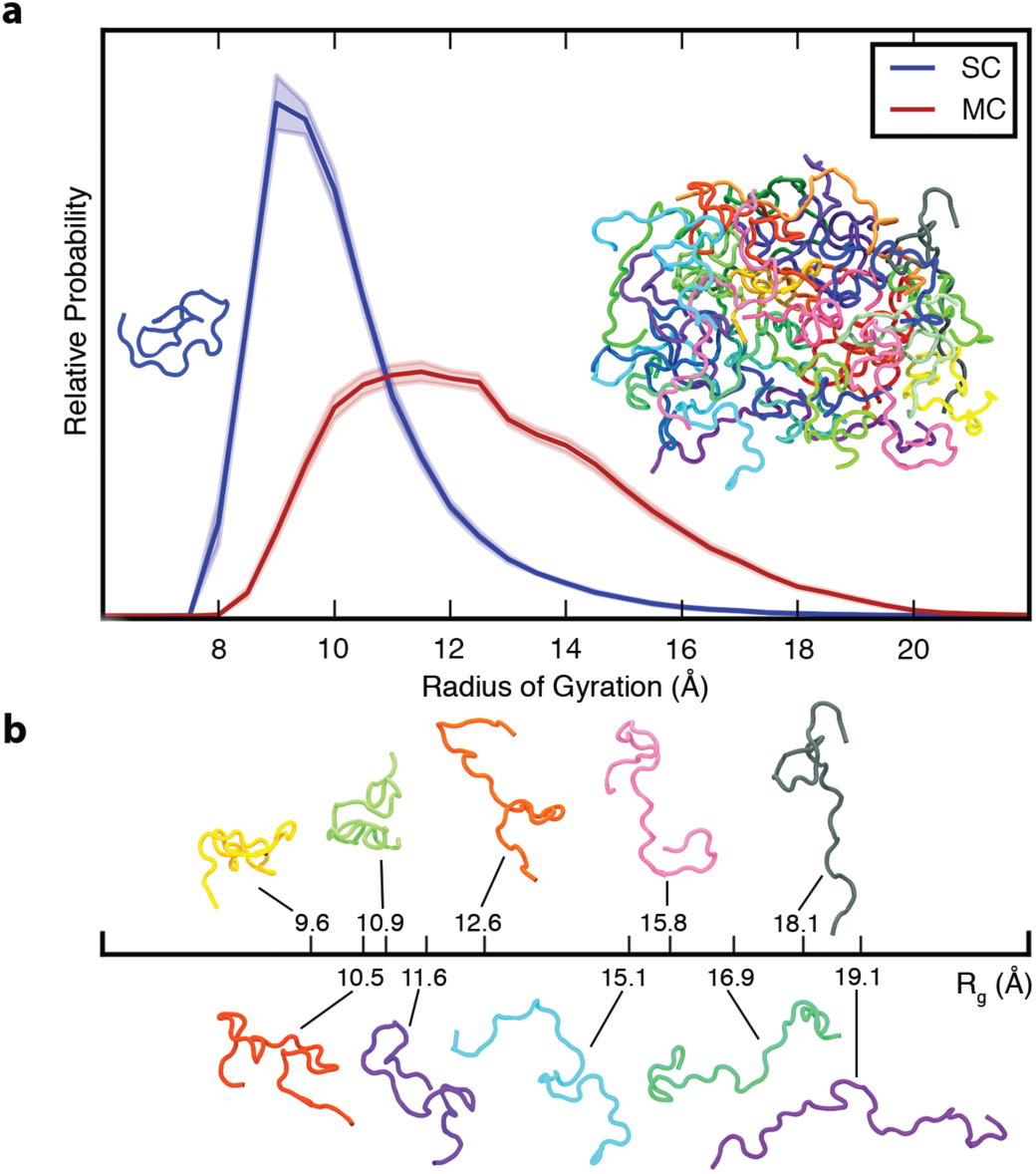
Ensemble-averaged polypeptide chain dimensions. **(a)** Probability distribution of the radius of gyration, R_g_, for SC (blue) and MC (red). Insets show representative backbone conformations of peptide monomer (left) and aggregate (right). **(b)** Aggregated chains are colored individually and ten of them are also shown below with their corresponding R_g_ to illustrate the conformational heterogeneity. In all figures, error bars indicate standard error. All the results reported in this and subsequent figures were obtained at 298 K unless otherwise noted.

High conformational disorder in the aggregate is corroborated by multiple, independent experimental observations on elastin: elastin fibres are optically isotropic;^34^ the backbone carbonyl order parameter is less than 0.1;^35^ carbon chemical shifts are consistent with random coil secondary structure;^35^ and neutron scattering experiments show that the polypeptide chains in elastin are highly mobile.^36^ High conformational entropy underpins the random network model of elastin structure and function initially proposed by Hoeve and Flory.^7,8^ In this model, conformational entropy decreases when the chains are stretched and increases upon relaxation, thereby driving elastic recoil. Consistent with the random network model, we find that the peptide chains in the aggregate are indeed highly disordered.

### Disordered, but not random: a probabilistic description of intra- and interchain interactions

In order to characterize the complex conformational landscape of the disordered ELP chains, we obtained a statistical picture of the different conformational states and interactions accessible to the chains in solution and in the aggregate. Statistical maps of the two types of peptide-peptide interactions, backbone hydrogen bonds and non-polar sidechain contacts, reflect highly-disordered conformational ensembles (Fig. 2). Secondary structure is sparse and limited to transient (sub-ns) hydrogen-bonded turns between residues close in sequence (near-diagonal elements in Fig. 2a, b). The most populated structures are VPGV and GVGV β-turns. The propensity of each of these local interactions, which peaks at 20%, is remarkably well conserved upon aggregation (Table 1), while the total number of peptide-peptide hydrogen bonds per chain increases only moderately (Fig. 2c). This evidence for fluctuating β-turns is consistent with spectroscopic data^1,20^ but not with the β-spiral, a highly-ordered structural model of elastin which requires that repeated β-turns be formed simultaneously,^6^ which is never observed in the simulations. As such, our results reconcile spectroscopic evidence for local secondary structure^1,20 37^ with global conformational disorder.

**Figure 2.**
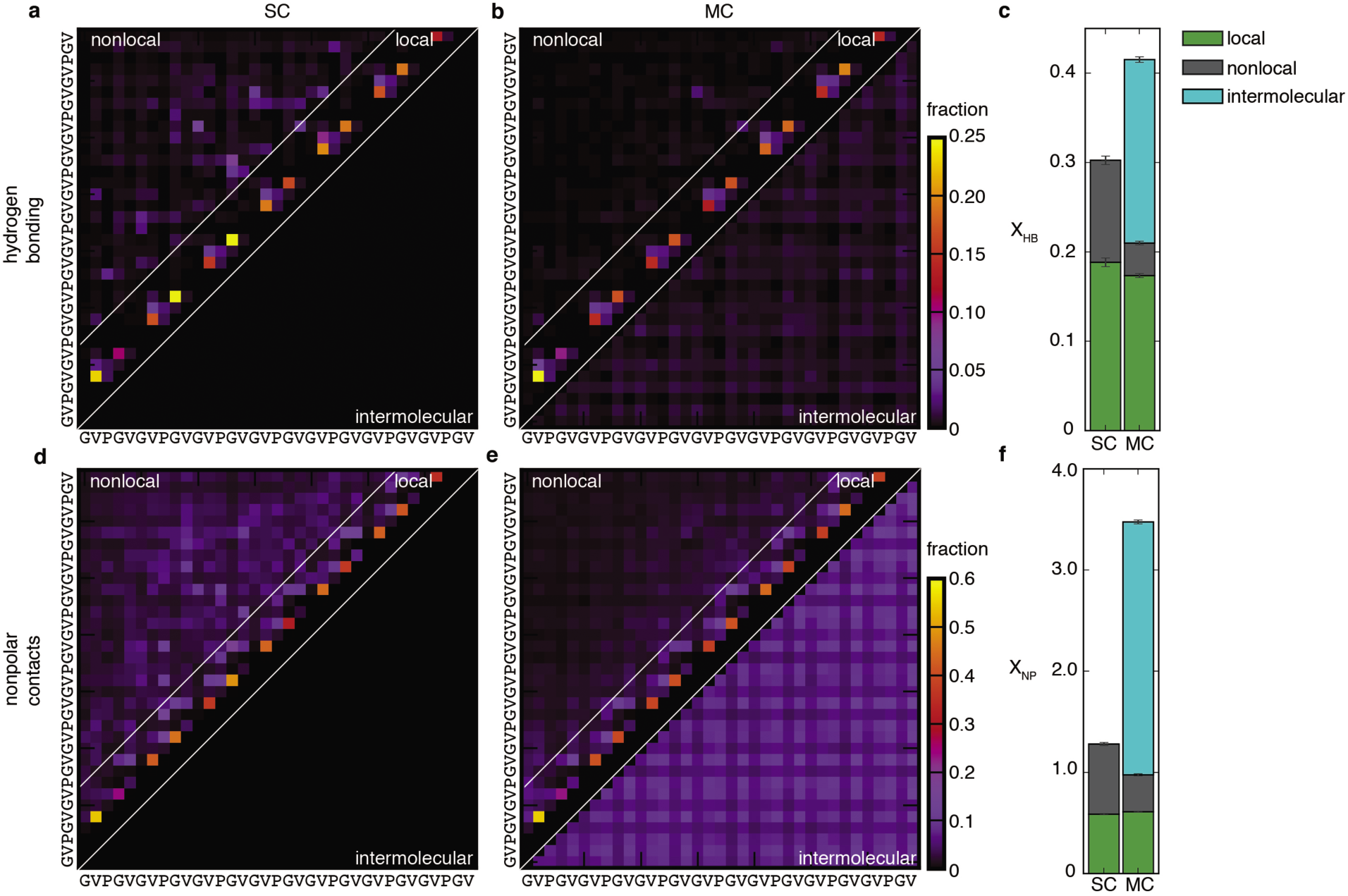
Peptide-peptide interactions. Probabilistic description of hydrogen-bonding (top row) and non-polar (bottom row) interactions of SC (**a** and **b**) and MC (**d** and **e)** systems. Panels (**(a), (b), (d)** and (**e)** are contact maps for pairwise interactions between residues. The color of each square indicates the fraction of conformations in the ensemble for which that interaction is present. Nearest- and next-nearest-neighbour contacts are excluded for clarity in **(b)** and **(e)**. Local interactions consist of sparse backbone hydrogen bonds and corresponding non-polar contacts. Nonlocal interactions consist primarily of non-specific non-polar contacts between sidechains. With the absence of preferred nonlocal interactions, the statistical picture of the conformational ensembles is remarkably simple. Upon aggregation, local structure propensities are retained as nonlocal hydrophobic contacts become intermolecular contacts (below the diagonal in **e**). **(c)** Average number of chain-chain hydrogen bonds per residue, X _HB_. **(f)** Average number of non-polar contacts per residue, X_NP_. Both X_HB_ and X_NP_ are the sum of intramolecular (local and non-local) and intermolecular contributions. Details of the structural analysis methods are provided in Supplementary Information.

**Table 1.** Populations and Lifetimes of Hydrogen Bonded Turns.

| Turn | Population (%) |  | Lifetime (ns) |  |
| --- | --- | --- | --- | --- |
|  | SC | MC | SC | MC |
| VPGV<sup>1</sup> | 18 ± 1 | 16 ± 2 | 0.18 ± 0.02 | 0.19 ± 0.01 |
| VPGVG | 5.9 ± 0.9 | 4.6 ± 0.7 | 0.9 ± 0.3 | 0.8 ± 0.1 |
| PGV | 1.5 ± 0.1 | 2.0 ± 0.2 | 0.008 ± 0.002 | 0.009 ± 0.001 |
| PGVG | 2.3 ± 0.2 | 2.9 ± 0.3 | 0.30 ± 0.05 | 0.39 ± 0.03 |
| GVGV | 20 ± 3 | 17 ± 2 | 0.15 ± 0.02 | 0.24 ± 0.07 |

<sup>1</sup>A hydrogen-bonded turn refers to a hydrogen bond between the first and last residue in the sequences shown. For example, the VPGV turn has a hydrogen bond between valine 1 and valine 4.

Even as it preserves local structural propensities, self-aggregation results in the replacement of nonlocal intra-molecular interactions by inter-molecular interactions (Fig. 2). In particular, the nonlocal non-polar contacts that characterize the collapsed isolated chain (Fig. 2d) give way to non-specific interactions with neighboring peptides (Fig. 2e). The average number of non-polar contacts per chain nearly triples upon self-assembly (Fig. 2f) with a commensurate decrease in the hydration of non-polar sidechains (Fig. 3b), indicating that the hydrophobic effect strongly contributes to the formation and the structure of the aggregate. Accordingly, the hydrophobic effect is the major driving force for elastic recoil in several earlier models of elastin. ^6,9–11,16^ Contrary to these models, however, significant hydrophobic burial is achieved even in the absence of a well-ordered structure.

**Figure 3.**
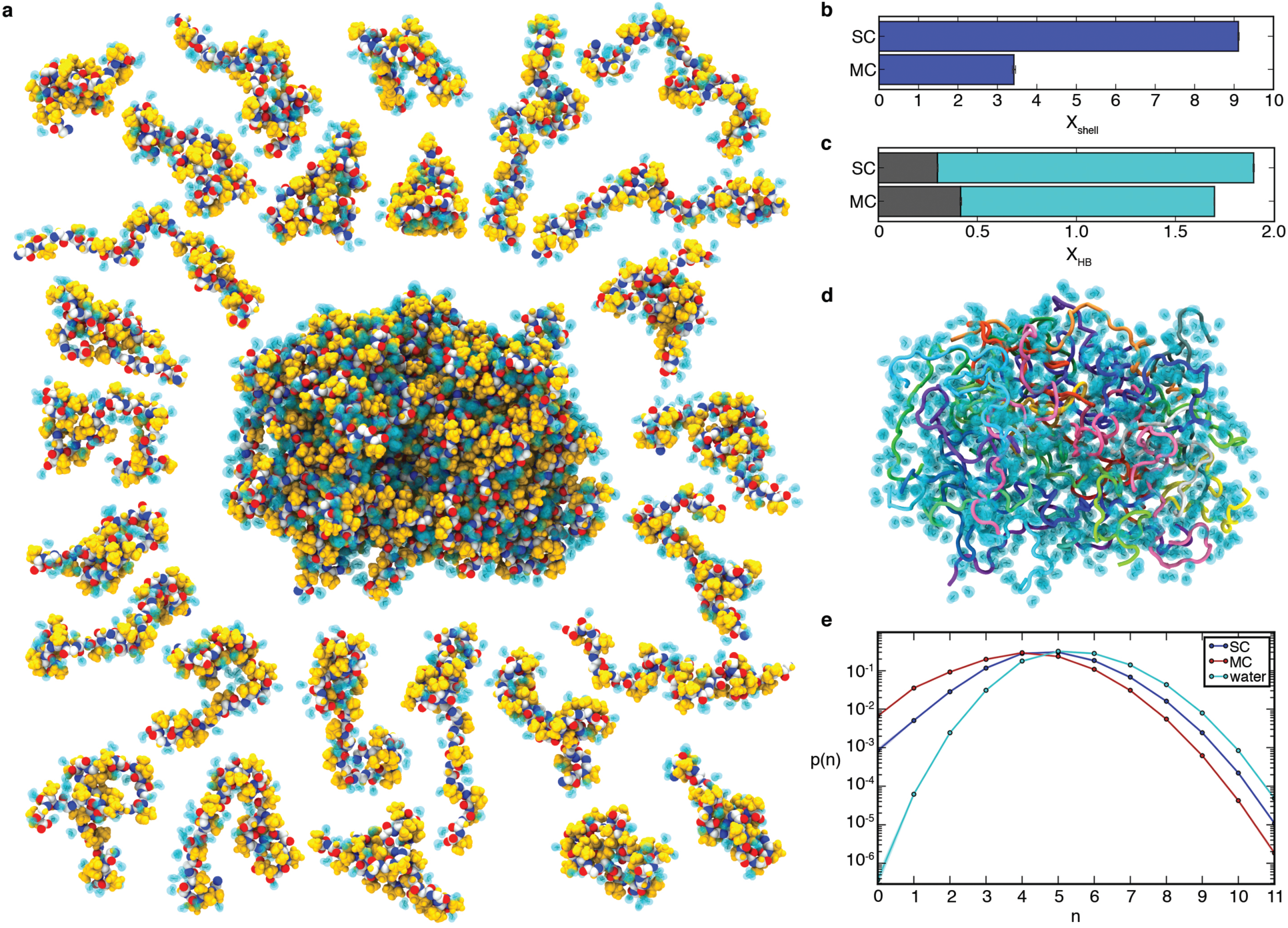
Peptide hydration in the liquid-like aggregate. **(a)** Representative conformation of the aggregate with non-polar side chains (yellow), peptide backbone (oxygen, red; carbon, white; nitrogen, blue), and hydrogen-bonded water molecules (cyan). Peptide chains are shown individually on the periphery with bound water molecules. **(b)** Average number of water molecules in the hydration shell per residue, X_shell_, for SC and MC systems (see Supplementary Information). **(c)** Average number of hydrogen bonds per residue, XHB. XHB is the sum of peptide-peptide hydrogen bonds (grey) and peptide-water hydrogen bonds (cyan), for both the SC and MC systems. **(d)** Same conformation as in panel a with bound water molecules shown as a transparent surface and peptides coloured individually. **(e)** Probability distribution, p(n), of water coordination number, n, for water molecules in the hydration shell of the SC (blue) and the MC (red) systems and in bulk water (cyan) at 298 K. Peptide-bound water molecules in the aggregate have fewer neighbors. The lines are shown to guide the eye.

### Hydration and disorder of the polypeptide backbone underlie elasticity

While self-assembly effectively buries non-polar side chains (Fig. 2e, f), disorder of the polypeptide backbone precludes the formation of a water-excluding hydrophobic core. Since there is only a moderate amount of secondary structure, a majority of backbone peptide groups do not form peptide-peptide hydrogen bonds (Fig. 2c). Instead, water molecules remain within the aggregate (Fig. 3a, d, e) in order to satisfy the hydrogen-bonding requirements of backbone groups (Fig. 3c). As a result, there is little loss of backbone hydration upon peptide self-assembly (Fig. 3c), even as the side chains become dehydrated (Fig. 3b). The high degree of hydration of the aggregate (40.0 ± 0.2% water content by mass, based on the number of water molecules hydrogen-bonded to the peptide) is consistent with experimental measurements of water content between 40 and 60 % for elastin derived from various species.^17^ In fact, the water content of both the monomer and the aggregate is so high that the probability of any five-residue segment to be completely dehydrated is essentially zero (Fig. S9b). This result clearly indicates that both systems lack a water-excluding hydrophobic core, a consequence of the high degree of conformational disorder and lack of extended secondary structure imparted by the high proline and glycine content.^28^

Maintaining both disorder and hydration upon aggregation is crucial for elastic recoil, but it is equally important that elastin and other self-assembled elastomeric proteins avoid the formation of the cross-β-structure characteristic of amyloid fibrils,^28^ which are postulated to be a thermodynamically stable state for any polypeptide chain under appropriate solution conditions. ^38^ Like the native state of globular proteins, the structure of amyloids is characterized by a water-excluding core and extensive backbone self-interactions. The liquid-like structure of elastin is incompatible with both protein folding and the formation of amyloid, and it is achieved through a high combined proportion of proline and glycine residues.^28^ Both proline, with its fixed ϕ dihedral angle and absence of amide hydrogen, and glycine, with its high entropic penalty for conformational confinement, inhibit the formation of α-helix and β-sheet structure, and serve to maintain a high degree of hydration and structural disorder by preventing the formation of a compact, water-excluding core. The essential role of proline and glycine in governing disorder, hydration, and elasticity was shown to extend to many other self-assembled elastomeric proteins,^28^ a finding corroborated by a recent study of an array of proline- and glycine-rich disordered proteins^39^ as well as by studies of various classes of spider silks.^40,41^ Taken together, these results point to a fundamental relationship between sequence composition, conformational disorder, and elastomeric properties of self-assembled elastomeric proteins, including elastin, spider silk, and resilin.

### Approaching the state of maximum conformational disorder: the liquid structure of elastin

The structural ensemble of the aggregate is disordered (Fig. 1), yet contains a significant propensity for secondary structure in the form of transient hydrogen-bonded turns (Fig. 2b, Table 1) and is highly hydrated (Fig. 3, Fig. S9). To understand why aggregation induces peptide expansion (Fig. 1a), we examine our results in terms of solvent quality. In a poor solvent, solute-solute interactions are energetically more favorable than solvent-solute interactions, leading to a collapse of the polymer chain. Inversely, solvent-solute interactions are preferred in a good solvent, leading to chain expansion. In the ideal limit between the two (the so-called “θ-solvent”), there is no preference for one type of interaction over the other. In this minimally-constrained state, the polymer becomes maximally disordered. The Flory theorem states that in the liquid, phase-separated state, the “polymer melt”, the polymer chains make extensive interactions with one another and become their own θ-solvent, since intramolecular and intermolecular interactions are chemically indistinguishable; as a result of which the chains reach a state of maximal disorder.^42–44^ Although disordered protein aggregates have been hypothesized to resemble polymer melts,^45,46^ whether or not they satisfy the Flory theorem is unknown.

The aggregated ELP chains are extensively solvated by one another (Fig. 2c, f), suggesting that they may approach the ideal, θ-solvent limit. To quantify the extent to which the chains approach this ideal limit, we compared the intra-chain distance scaling of the conformational ensembles in the simulations to the scaling expected for the ideal, θ-solvent limit (Fig. 4). As a model for this ideal state, we use the residue-specific model for a random-coil polypeptide developed by Flory and co-workers.^43,47^ In their model, rotation about the backbone dihedral angles, ϕ and ψ, is treated as independent of the conformation of neighboring residues in the chain. We update their method to include residue-specific transformation matrices derived directly from the simulation data (see the detailed description of the method in Supplementary Information). The internal distance scaling profile for the Flory model of the ideal state is fit by a power law with exponent α = 0.54, which is very similar to the expected dimensions of an ideal Flory random coil homopolymer, for which α = 0.5.^43^ Similarly, the dimensions of the chains in the aggregate, with an exponent α = 0.46, closely approach the ideal limit as well. In contrast, the dimensions of the ensemble in solution, with an exponent of α = 0.28, are consistent with those of a polymer in a poor solvent, for which α = 1/3. This result is expected, given the highly hydrophobic composition of the ELP sequence, and the fact that water is a poor solvent for hydrophobic residues. The internal distance scaling profile of the chain in solution exhibits a slight upturn at large sequence separations. This deviation from the behavior expected for a perfectly collapsed, globular homopolymer likely arises from the presence of seven proline residues in the sequence, which induces a local stiffness that significantly limits the conformations accessible to the polypeptide chain. Furthermore, the conformational landscape of the ELP in solution is more complex than that of a simple homopolymer in a poor solvent due to the presence of specific, highly-populated turns (Table 1 and Fig. 2).

**Figure 4.**
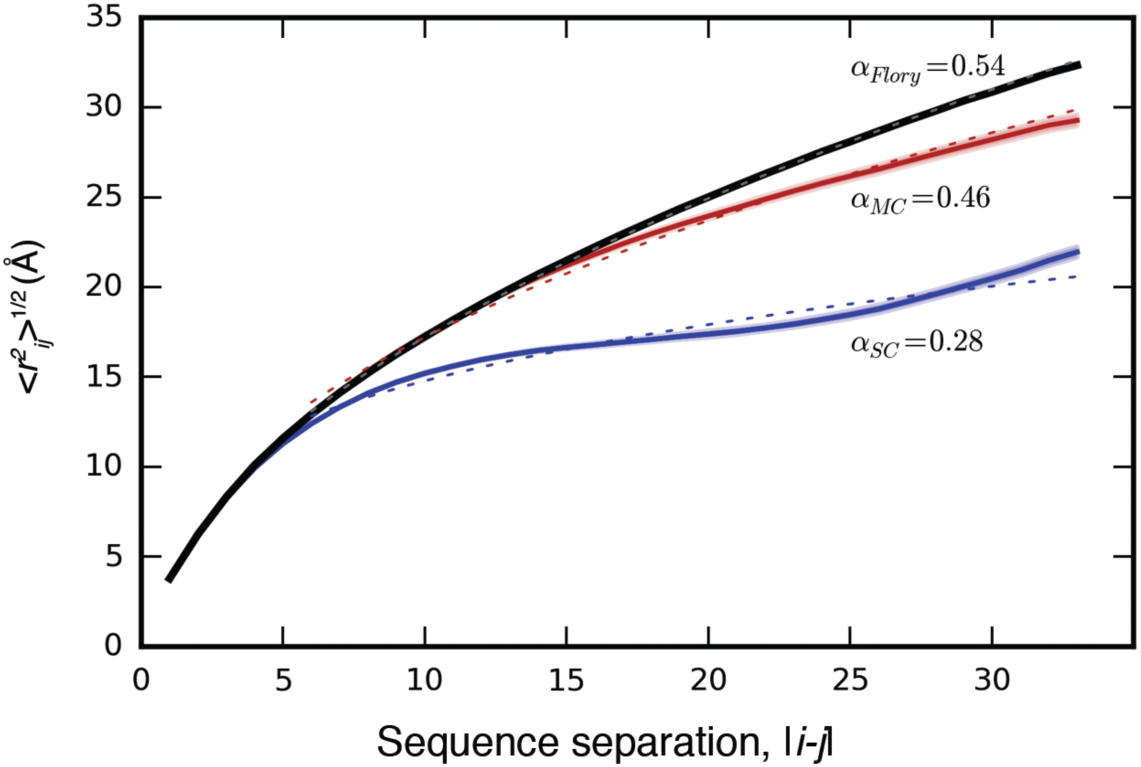
Intrachain Distance Scaling and Comparison to the Ideal, Random-Coil State. Root-mean-square distance, <*r_ij_*^2^>^1/2^, between residues *i* and *j* as a function of sequence separation, |*i*-*j*|, for SC (blue) and MC (red), and for the ideal, random coil state modeled using the SC simulations according to the method of Flory et al. (black)^43 47^, described in detail in Supplementary Information. Shading indicates standard error. In each case, the dotted line indicates the power law fit to the data, with the exponent α provided next to each curve. Here, the fits were carried out for |i-j|> 5 because the length of a blob-sized segment was found to be 5 residues (Figure S9), and the distance scaling within a blob differs from that outside of a blob.

The dimensions of the chains in the ELP aggregate approach the dimensions predicted for maximally-disordered chains (Fig. 4), and therefore the dimensions expected in a polymer melt. Deviation from ideality reflects the finite size of the aggregate, finite chain length, the presence of local secondary structure (Fig. 2c), and persistent hydration (Fig. 3, Fig. S9). These results suggest that elastin—and polypeptide chains in general — cannot make polymer melts in the idealized, solvent-excluding sense because backbone groups must form hydrogen bonds either with each other, which leads to ordering, or with water molecules, whose presence is required for disorder. Instead, the chains adopt a disordered state with significant backbone hydration, as seen in the representative structure shown in Fig. 3a, d. The fact that the dimensions of chains within the aggregate are much closer to the ideal state than the single chain in solution indicates that conformational entropy increases dramatically upon aggregation and is an important driving force for aggregation. Taken together, these findings demonstrate how a protein aggregate may approach maximal conformational disorder.

### Peptide chain dynamics in solution and in the aggregate

The conformational ensembles of the peptide in solution and in the aggregate differ significantly with respect to chain dimensions (Fig. 1, 4), long-range contacts, hydrophobic interactions (Fig.2), and hydration (Fig. 3). Despite these large global structural differences, the ensembles strongly resemble each other in terms of local secondary structure: the populations and lifetimes of the hydrogen-bonded turns are nearly identical for both SC and MC systems (Table 1). While the dynamics of turn formation are similar, non-local dynamics of the chain differs dramatically in solution and in the aggregate. In particular, the lifetime of the open state (defined as the N- and C-termini not being in contact) increases more than fifty times upon aggregation (21 ± 1 ns vs. 1140 ± 30 ns in the MC and SC systems, respectively; Fig. S6).

Reptation theory predicts characteristic signatures for the dynamics of polymer chains in melts:^48^ short timescale motions are predicted to obey Rouse-like dynamics, whereas long timescale motions should be strongly affected by the confinement imposed by neighbouring chains. To determine whether the dynamics of aggregated elastin peptides is characteristic of a melt, we analyzed the diffusion of the central residue of each chain (Fig. S10). This residue exhibits anomalous diffusion (or sub-diffusion), with its mean-square displacement obeying a power law with exponent α = 0.578. This value is similar to the exponent close to 0.6 found by Harmandaris *et al.* in atomistic simulations of polyethylene melts^49^ and is intermediate between 1/2 and 2/3, as expected respectively for a Rouse chain^50,51^ and for the Zimm model, an extension of the Rouse model that accounts for hydrodynamic interactions between chain monomers.^51,52^ Importantly, we find no crossover to a regime with α = 1/4, as one would expect if the chain were moving as if confined in a tube formed by the neighboring chains (that is, the entangled polymer melt regime described by de Gennes^48^). The chain length used here may be too short to observe this crossover. Consistent with this hypothesis, Ramos *et al.^53^* found no crossover for the shortest chain length of hydrogenated polybutadiene that they studied (36 monomers compared to 35 residues in the chain studied here). The absence of crossover may also be due to insufficient simulation length, or to the fact that the chains remain highly hydrated in the aggregate and are not characteristic of a solvent-excluding melt.

## CONCLUSIONS

The present study provides the first atomistic description of a melt-like, disordered protein state, which may be called the liquid state of proteins. The model of elastin-like aggregates derived here represents the first detailed model of a protein coacervate. In spite of its moderate size, this molecular system emulates a biphasic liquid in which both phases have attained a state of higher entropy than in the mixed system: the higher entropy of the aqueous phase results from the hydrophobic effect, which is manifested by a three-fold increase in burial of non-polar groups for each polypeptide chain in the aggregate compared to the monomeric form. In the aggregate, the polypeptide chain approaches a state of maximal conformational entropy as predicted by the Flory theorem. As such, the above results show how a classic concept of polymer theory, the polymer melt, is realized in an important but poorly-understood structural protein, and demonstrate the relevance of this concept to the self-assembly (coacervation) and mechanical properties of elastin.

Our results support a unified model of elastin structure and function that recapitulates experimental data and reconciles key aspects of previous qualitative models. The biological function of elastin is incompatible with a unique, ordered structure. In the functional state, its hydrophobic domains form a water-swollen, disordered aggregate characterized by an ensemble of many degenerate conformations with significant backbone hydration and fluctuating local secondary structure. The combination of two entropic driving forces, the hydrophobic effect and polypeptide chain entropy, governs both self-assembly and elastic recoil. These effects are intimately linked. On the one hand, replacing intramolecular hydrophobic contacts by intermolecular ones allows the aggregated chains to approach a state of maximal conformational entropy, which underlies the elastic recoil central to elastin’s function. On the other hand, the association of non-polar groups and the formation of a maximally-disordered, liquid-like protein aggregate also drive elastin’s liquid-liquid phase separation.

Remarkably, our findings defy conventional wisdom about protein folding, aggregation, and disorder: (1) although the structure of elastin-like aggregates is nearly maximally disordered, it is not random but instead contains well-defined and significantly-populated secondary structure elements in the form of hydrogen-bonded turns; (2) however, because turns are local, sparse, and transient, the polypeptide backbone remains significantly hydrated overall; as a result, (3) the hydrophobic side-chains cannot form a compact, water-excluding core even though they are significantly buried.

The detailed model of the liquid phase of proteins obtained here for elastin is of direct relevance to the self-assembly and mechanical properties of other self-assembled elastomeric proteins, with which elastin shares a high content in proline and glycine. ^28,39^ In addition, by revealing the structural and physico-chemical basis for the phase separation of elastin, this study also provides a frame of reference for understanding the phase separation of other functional disordered proteins, including the FG-nucleoporins that compose the selectivity barrier of the nuclear pore complex,^54^ and low-complexity protein assemblies implicated in the intracellular phase separation of membraneless organelles.^55-60^

## METHODS

Atomistic MD simulations with explicit water were performed on an elastin-like peptide (ELP), (GVPGV)_7_, successively as an isolated chain (single chain, SC) and an aggregate of twentyseven chains (multi-chain, MC). This sequence, derived from a hydrophobic domain of chicken elastin, is the most extensively studied elastin repeat motif.^1^ An accumulated simulation time of over 200 *μ*s was required to reach statistical convergence. A full description of simulation methods and structural analysis is provided in Supplementary Information.

## Acknowledgements

We thank F.W. Keeley, L. Muiznieks, R.V Pappu, S.G. Whittington, and V. M. Burger for discussions. Computations were performed on the GPC supercomputer at the SciNet High-Performance Computing Consortium and on the supercomputer Guillimin from McGill University. SciNet is funded by the Canada Foundation for Innovation under the auspices of Compute Canada; the Government of Ontario; Ontario Research Fund - Research Excellence; and the University of Toronto. Guillimin is managed by Calcul Québec and Compute Canada; its operation is funded by the Canada Foundation for Innovation, NanoQuébec, RMGA and the Fonds de recherche du Québec - Nature et technologies. This work was supported by Canadian Institutes for Health Research operating grant MOP84496 to R.P. and by a Canada Graduate Scholarship from the Natural Sciences and Engineering Research Council and a Scholarship from the Research Training Centre at the Hospital for Sick Children to S.R.

## Author Contributions

S.R. and R.P. designed the study; S.R. carried out the research and analysed data; S.R. and R.P. wrote the paper.

